# Multiple optic gland signaling pathways implicated in octopus maternal behaviors and death

**DOI:** 10.1101/340984

**Authors:** Z Yan Wang, Clifton W Ragsdale

## Abstract

Octopus optic glands employ a multiplex progression of signaling molecules to regulate maternal behaviors.

**1. Abstract:** Post-reproductive life in the female octopus is characterized by an extreme pattern of maternal care: the mother cares for her clutch of eggs without feeding until her death. These maternal behaviors are completely eradicated if the optic glands, the octopus analog of the vertebrate pituitary gland, are removed from brooding females. Despite the optic gland’s importance in regulating maternal behavior, the molecular features underlying optic gland function are unknown. Here, we identify major signaling systems of the *Octopus bimaculoides* optic gland. Through behavioral analyses and transcriptome sequencing, we report that the optic gland undergoes remarkable molecular changes that coincide with transitions between behavioral stages. These include the dramatic up- and down-regulation of catecholamine, steroid, insulin, and feeding peptide pathways. Transcriptome analyses in other tissues demonstrate that these molecular changes are not generalized markers of aging and senescence, but instead, specific features of the optic glands. Our results provide strong evidence for the functional conservation of signaling molecules across evolutionarily distant species. For example, elevated levels of insulin growth factor binding proteins are associated with cachexia-like tissue wasting in flies, humans, and, reported here, octopuses. Our study expands the classic optic gland-pituitary gland analogy and indicates that, rather than a single “self-destruct” hormone, the maternal optic glands employ multiple pathways as systemic hormonal signals of behavioral control.

## 2. Introduction

Octopuses and other soft-bodied (coleoid) cephalopods are short-lived, semelparous animals: adults die after a single reproductive bout (Rocha et al., 2001). Typically, octopuses lead solitary lives and come together only to mate (Wells, 1978). Females then store sperm in specialized compartments in their oviducal gland (Wells, 1978). As eggs are laid, they pass through the oviducal gland and are fertilized. The female octopus anchors her eggs to substrate with mucal secretions and tends to her clutch as the embryos develop. During this brood period, she rarely leaves her clutch and abstains from food. By the time of hatchling, the female dies (Anderson et al., 2002; Hanlon and Messenger, 1998; Wells, 1978).

In a key experiment from 1977, Jeremy Wodinsky surgically resected optic glands from female octopuses that were brooding their clutch of eggs and had stopped feeding. Removal of the optic glands led to substantial behavioral changes: females abandoned their clutches, resumed feeding, gained weight, and some even mated again. Glandectomized individuals lived 5.75 months longer than their intact counterparts, leading Wodinsky to conclude that the optic gland and optic gland secretions constituted an octopus “self-destruct system” (Wodinsky, 1977). The molecular features underlying optic gland signaling have not been explored with modern investigative techniques and the putative “optic gland hormone” (Wells, 1978) remains unidentified to this day.

Classic work from Wells and Wells (1959) established that the optic glands are also necessary for the proper timing of sexual maturation. The optic glands are situated on the optic stalks, nestled between the large kidney-shaped optic lobes and the central brain. They are known to receive inhibitory signals from the subpedunculate lobe of the supraesophageal brain (O’Dor and Wells, 1978; Wells and Wells, 1959). At sexual maturity, this inhibition is released; the optic gland swells in size and darkens in color (Wells and Wells, 1975). This change causes the gonads and reproductive organs to mature (Wells and Wells, 1959). Wells posited that the optic glands are the octopus analog to the vertebrate pituitary gland, acting as intermediaries between brain control centers and peripheral target organs (Wells and Wells, 1969).

To gain insight on the molecular features underlying post-reproductive behavioral changes, we sequenced optic gland transcriptomes of the California two-spot octopus, *Octopus bimaculoides*. *O. bimaculoides* lives off the coast of southern California and northern Mexico. We could observe normal maternal behaviors of this species in a laboratory setting. These behavioral findings guided our optic gland RNA sequencing and interpretation of bioinformatics results.

## 3. Materials and Methods

### 3.1 Animals and animal facilities

Wild-caught mated and non-mated female California two-spot octopuses (*O. bimaculoides*) were purchased from Aquatic Research Consultants (Catalina Island, CA) and shipped overnight to Chicago, IL.

All was performed in compliance with the EU Directive 2010/63/EU guidelines on cephalopod use and the University of Chicago Animal Care and Use Committee (Fiorito et al., 2014; Fiorito et al., 2015; Lopes et al., 2017). Mated females were kept in their home dens with clutches as much as possible. Clutches were sparingly separated from females for other experiments. Animals were individually housed in artificial seawater (Tropic Marin) in 20- or 50-gallon aquaria, and offered a daily diet of live fiddler crabs (*Uca pugnax*, Northeast Brine Shrimp, Oak Hill, FL), cherrystone clam meat, and grass shrimp. Water temperature was maintained between 17-21°C and ambient room temperature at 21-23°C. Animal rooms were kept on a 12:12 hour light/dark cycle.

### 3.2 Behavioral Categorization

Upon arrival at our animal facilities, we examined octopuses for signs of injury such as pre-existing skin lesions and missing arm tips. These animals were excluded from further study. Octopuses were kept undisturbed for 4 days so they could habituate to our aquarium setup and recover from the distress of flying. On the 4^th^ day, we began offering live prey items. Each day, females had 4 live fiddler crabs available to them. Animals were observed twice daily (AM and PM) and recorded overnight with a Sony Handycam FDR-AX100 to characterize their behaviors. We observed 20 non-mated animals for at least 4 days before sacrificing and sexing them (see below). 11/20 non-mated animals were males and were excluded from the study. In addition, we longitudinally observed 16 brooding females and categorized them into behavioral stages based on the traits detailed in the Results section and summarized in Table 1.

**Table 1.**
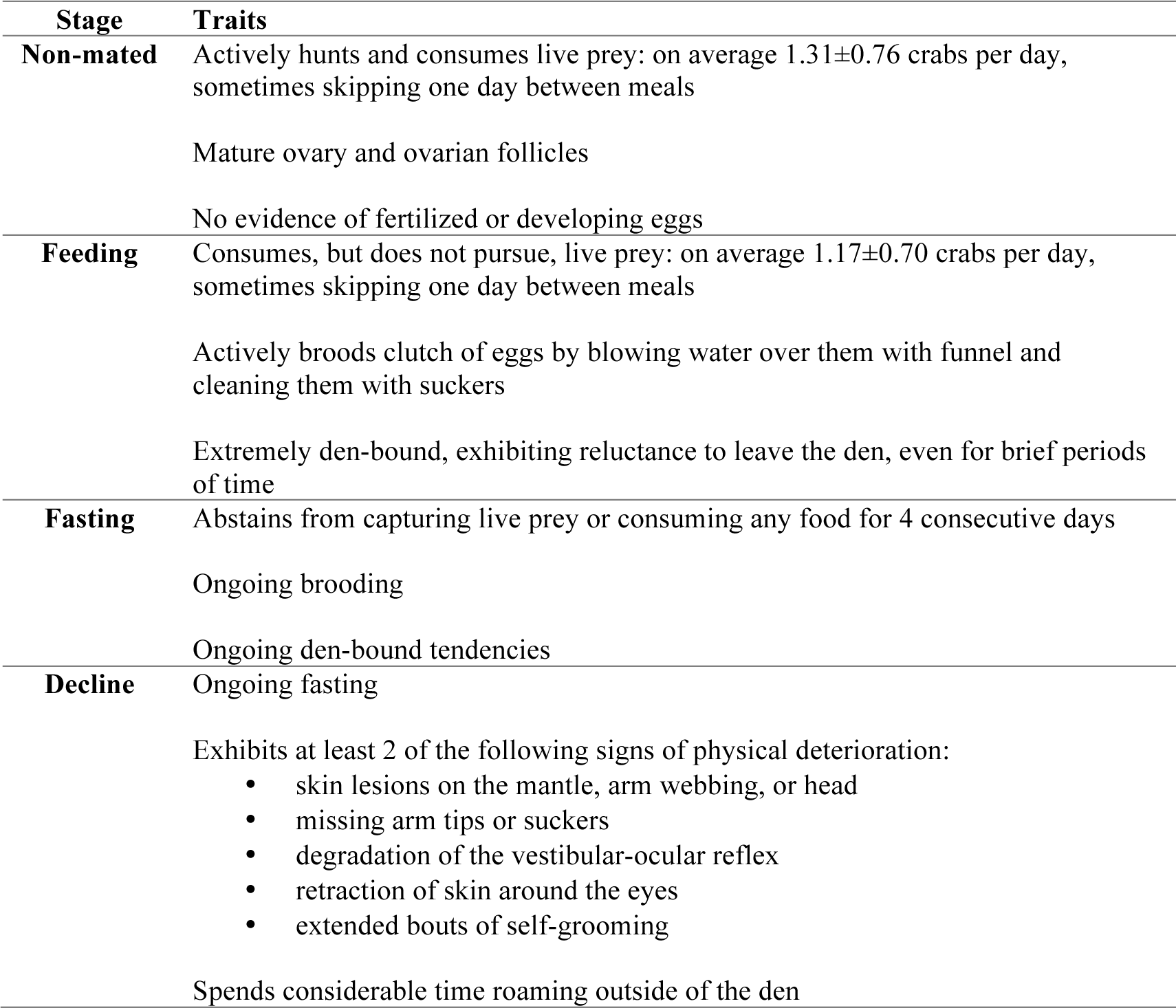
Octopus female reproductive life is characterized by four stages of behavior.

| Stage | Traits |
| --- | --- |
| Non-mated | Actively hunts and consumes live prey: on average 1.31±0.76 crabs per day, sometimes skipping one day between meals<br>Mature ovary and ovarian follicles<br>No evidence of fertilized or developing eggs |
| Feeding | Consumes, but does not pursue, live prey: on average 1.17±0.70 crabs per day, sometimes skipping one day between meals<br>Actively broods clutch of eggs by blowing water over them with funnel and cleaning them with suckers<br>Extremely den-bound, exhibiting reluctance to leave the den, even for brief periods of time |
| Fasting | Abstains from capturing live prey or consuming any food for 4 consecutive days<br>Ongoing brooding<br>Ongoing den-bound tendencies |
| Decline | Ongoing fasting<br>Exhibits at least 2 of the following signs of physical deterioration:<br>• skin lesions on the mantle, arm webbing, or head<br>• missing arm tips or suckers<br>• degradation of the vestibular-ocular reflex<br>• retraction of skin around the eyes<br>• extended bouts of self-grooming<br>Spends considerable time roaming outside of the den |

### 3.3 Tissue collection

Octopuses were submerged in 5-10% ethanol/seawater for at least 20 minutes to achieve deep anesthesia, then decerebrated (mantle and arms removed from the head) (Butler-Struben et al., 2018; Gleadall, 2013). The head was dissected on ice in diethyl pyrocarbonate-treated phosphate buffered solution (DEPC-PBS). Optic glands were accessed between the eyes, harvested, flash frozen in Trizol (Invitrogen), and stored at -76°C until RNA extraction. Because sexual dimorphism is difficult to observe in our species, animals were definitively sexed after tissue harvest. Only females with mature ovaries, ovarian follicles, and no evidence of fertilized eggs were considered non-mated females.

We also harvested arm tissue from the same cohort of animals. Transverse arm slices corresponding to the 10^th^ row of suckers distal from the mouth on the second arms on the right and left were cut into quadrants to facilitate tissue homogenization. Each arm piece was separately flash frozen in Trizol and stored at -76**°**C until RNA extraction.

### 3.4 RNA extraction and sequencing

Whole optic glands were homogenized with pellet pestles (Fisherbrand) in microcentrifuge tubes. Arm sections were homogenized in a Potter-Elvehjem tissue grinder (Kimble, Rockwood, TN). RNA was extracted using Trizol and PhaseMaker Tubes (Life Technologies Corporation, Carlsbad, CA) following manufacturer’s instructions. RNA integrity was checked with a Bioanalyzer 2100 (Agilent). Only samples with clean cephalopod rRNA peaks and little to no evidence of degradation were used. Tissue from the right arm was processed for RNA extraction and sequencing. The left arm was used in two samples of feeding females when RNA quality from the right arm was poor.

Total RNA was polyA-selected and directionally sequenced at the University of Chicago Genomics Facility on an Illumina HiSeq2500 machine, generating 100bp directional paired-end reads with an insert size of 300bp.

### 3.5 *de novo* transcriptome assembly and differential gene expression analyses

Following removal of adapters and low-quality sequences, reads were assembled with the Trinity software platform (v2.4.0) (Haas et al., 2013). Both individual and pooled transcriptomes were created (Table 2). Gene and isoform expression levels were estimated with RSEM (v1.2.31) (Li and Dewey, 2011) and differential expression was analyzed with edgeR (v3.7) (Robinson et al., 2009). Heatmaps were created in R with the heatmaps.2 function and colored with palettes from the RColorBrewer package.

**Table 2.** *O. bimaculoides* optic gland transcriptome assembly summary. Whole optic glands and transverse arm sections were harvested (see Methods). Separate libraries were prepared for each sample, and reads were later concatenated for transcriptome assembly. Statistics are based on transcript contigs. Total Trinity ‘genes’ refers to the number of transcript clusters generated by the assembly (isogroups), while Total Trinity Transcripts indicates the number of isoforms (isotigs). The Nx length statistic indicates that at least x% of the assembled transcript bases are found on contigs that are of at least Nx length; for example, at least 10% of the assembled bases in the unmated female optic gland transcriptome are found on transcript contigs that are at least 4012 bp long.

| Transcriptome | # of<br>libraries<br>prepared | Total<br>Trinity<br>'Genes' | Total<br>Trinity<br>Transcripts | N10<br>(bp) | N50<br>(bp) | Average<br>Contig<br>Length (bp) | Total<br>Assembled<br>Bases (bp) |
| --- | --- | --- | --- | --- | --- | --- | --- |
| Unmated Female Optic Gland | 3 | 267380 | 314115 | 4012 | 882 | 604.80 | 189975279 |
| Feeding Mom Optic Gland | 3 | 290774 | 344673 | 4078 | 909 | 616.68 | 212552829 |
| Fasting Mom Optic Gland | 3 | 254893 | 301599 | 4011 | 938 | 627.42 | 189228407 |
| Decline Mom Optic Gland | 2 | 217478 | 261725 | 4554 | 1165 | 688.47 | 180189329 |
| **Combined Optic Gland** | -- | 598083 | 706186 | 3897 | 767 | 574.65 | 405807628 |
| Unmated Female Arm | 3 | 120017 | 189097 | 5771 | 1781 | 904.15 | 170971875 |
| Feeding Mom Arm | 3 | 296853 | 406986 | 5196 | 1165 | 676.91 | 275491458 |
| Fasting Mom Arm | 3 | 183259 | 278194 | 5683 | 1621 | 823.35 | 229050350 |
| Decline Mom Arm | 2 | 153752 | 233040 | 5514 | 1650 | 841.44 | 196090319 |
| **Combined Arm** | -- | 493537 | 686453 | 4485 | 959 | 639.22 | 438793048 |

In the optic gland transcriptomes, 1,198 Trinity transcripts showed at least a 4-fold change between any two of the behavioral stages, at a p-value of 0.001. A phylogeny of differentially expressed genes was created by Euclidean clustering (Supplemental Fig 1). The tree was cut at a height of 0.3, which identified 25 subclusters. 20 genes from subclusters of interest (see below) were confirmed by PCR amplification.

## 4. Results

### 4.1 Adult life in the female O. bimaculoides can be divided into 4 behavioral stages

Sexually mature female octopuses are known to be active predators (Hanlon and Messenger, 1998). We confirmed this behavior in non-mated females in the lab: these animals spent time outside of the den (Fig 1A) and primarily hunted through a visually-directed jet-propelled “pounce” (Fig 1B and Movie S1) (Wells, 1978). The female first watches the prey out of one eye, then bobs her head up and down, likely acquiring depth information. Even while moving her head, her pupil is kept perpendicular to gravity (Fig 1A). Following a swift pounce, prey is captured in the interbrachial web (Fig 1B and C). Non-mated females consumed 1.31±0.76 crabs a day, sometimes skipping a day between meals.

**Fig 1.**
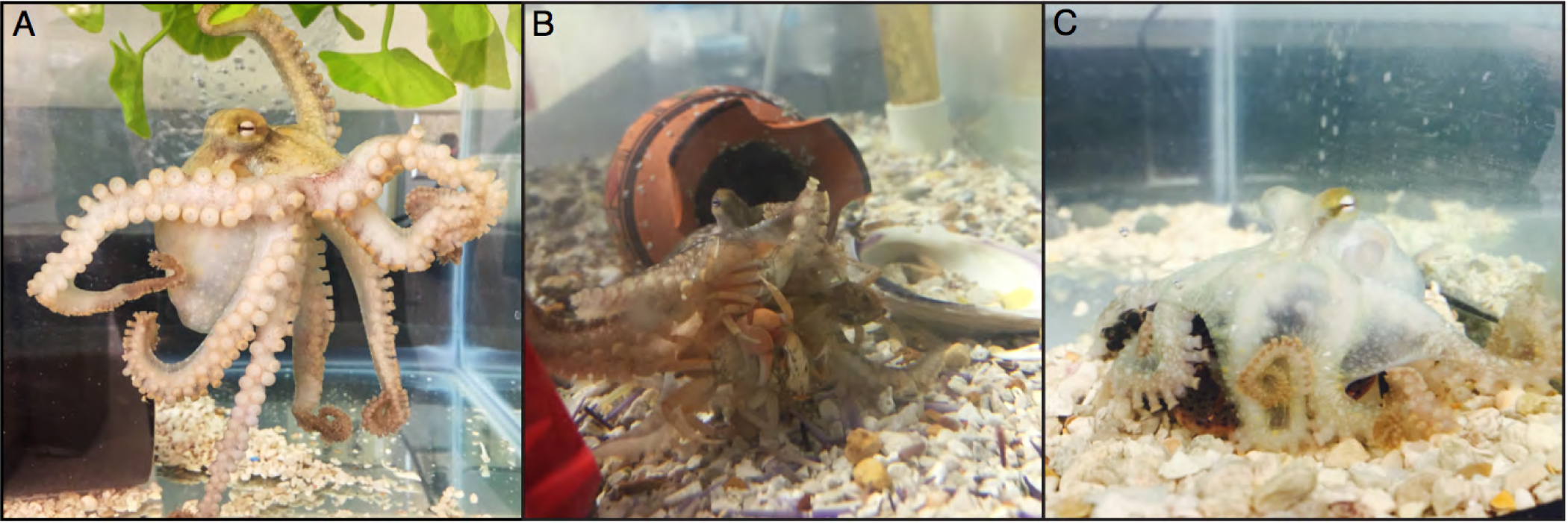
Non-mated, sexually mature females are active predators that hunt live prey. A) Females in this stage spend time outside of their home den. B) Non-mated females use visual direction to strike prey with a “pounce,” viewed here from the front. C) The webbing between the arms is spread wide to capture prey. Several fiddler crabs may be restrained under the interbrachial web at one time.

Mated females in the first stage of brooding actively tended their clutch by guarding their den, stroking eggs with their suckers, and blowing water over the eggs with their funnels (Hanlon and Messenger, 1998; Wells, 1978). In the lab, feeding mated females consumed 1.17±0.70 crabs per day, with fewer than 3 days without a meal. Feeding females, however, rarely left their egg clutches. Instead of capturing prey by stalking and pouncing, feeding females peered out of their dens with one eye and grabbed prey as they moved passed the mouth of the den (Fig 2A and B and Movie S1). The presence of empty crab carapaces provided additional confirmation that feeding females consumed food, rather than merely capturing and killing prey. On average, mothers fed for 8±2.71 days before starting to fast.

**Fig 2.**
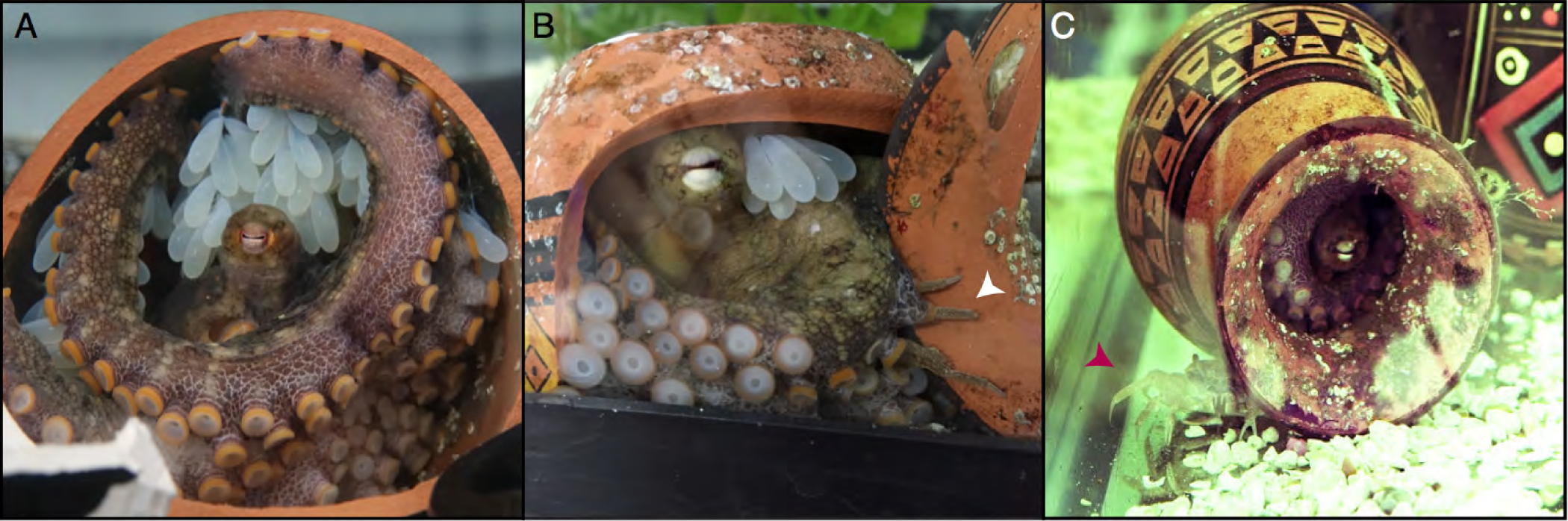
Brooding females feed and then fast. A) In the first stage of brooding, females actively tend to their eggs and feed. The pupils of feeding females remain perpendicular to gravity. B) Feeding females capture and kill live prey, but only without leaving the home den. White arrowhead indicates the legs of a captured fiddler crab extending from beneath the arms. C) Fasting females continue tending to their clutch but abstain from feeding even when live prey (red arrowhead) are within reach.

Females in the second stage of brooding continued to care for eggs but abstained from feeding, even though live prey items were available at all times (Fig 2C and Movie S1). Females who stopped feeding for 4 consecutive days were never observed to feed again, and we grouped these animals into the fasting stage.

After 11±8.49 days, fasting mothers began a rapid decline. Behavioral signs of decline, which have not been described previously, were observed before physiological signs (Fig 3A): females left their clutches and spent time outside of the den, languidly sitting on the bottom of the tank or persistently smashing themselves into the corners of the tank. The latter action led to the rapid formation of deep avulsions on the mantle or arms that did not heal (Fig 3B and C). Females also engaged in excessive self-grooming behaviors. Normally, octopuses run their first pair of arms over their mantles and head to remove ectoparasites and debris (Mather and Alupay, 2016). Decline mothers moved all of their arms to groom, but often arms would only pass over other arms, creating a turbulent mass of entangled limbs (Fig 3A). In a few cases, this over-grooming behavior was directly followed by self-cannibalization of the arm tips or suckers (Fig 3B and C and Movie S1). Physiological features of females in decline included a retraction of the skin around the eyes, general paling of skin color, and loss of motor tone (Anderson et al., 2002). In addition, the pupils also gradually ceased to remain perpendicular to gravity, raising the possibility that the central systems responsible for the vestibulo-ocular reflex deteriorate during this stage.

**Fig 3.**
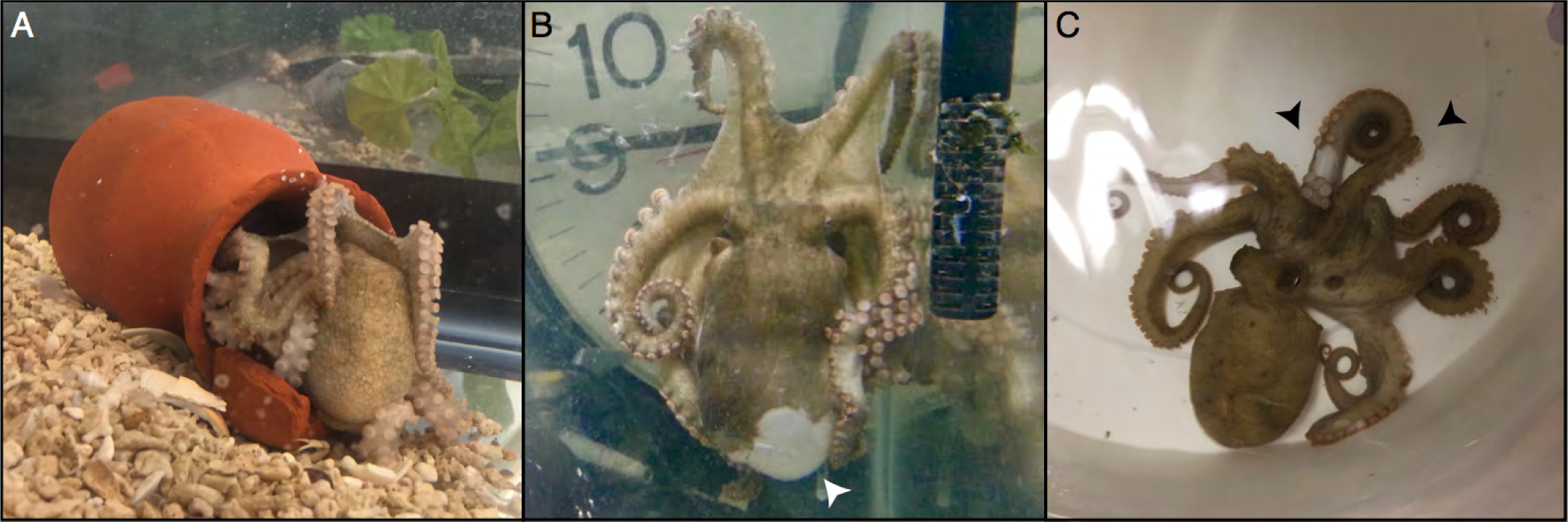
Females in decline continue fasting and undergo rapid senescence. A) Behavioral signs of decline include spending time outside of the home den and recruiting all the arms in bouts of aberrant, grooming-like behavior. Females also show physiological signs of decline, such as an apparent loss of muscle tone and unhealed wounds. B) Female with a patch of missing skin at the distal end of her mantle (white arrowhead). C) Female with several rows of missing suckers and a missing arm tip (black arrowheads), presumably from self-cannibalization.

In all cases, mated females progressed from feeding to fasting to decline before dying: that is, mated females did not transition from feeding directly to decline without fasting, and females did not die without showing indications of decline. Our observations suggest that after mating, female octopuses undergo a series of near-requisite behavioral transitions before death.

### 4.2 Transcriptomes of brooding females implicate many signaling systems in optic gland function

We assembled transcriptomes of left and right optic glands harvested from multiple individuals in each of the 4 stages (Table 2). Our assembled transcriptomes showed that the optic glands from non-mated animals were the most different from the other optic gland samples (Supplement Fig 2). After estimating transcript abundance with RSEM, we identified 1,198 differentially expressed transcripts with edgeR and grouped them into 25 subclusters (Supplement Fig 1). Wodinsky’s work suggested that optic gland signaling changes steadily over time until reaching a threshold that triggers fasting and later death (Wodinsky, 1977). In selecting subclusters for further study, we were guided by two criteria. We excluded 12 subclusters that showed a departure from monotonicity: for example, when expression peaked in only the feeding or fasting stages (Supplement Fig 3). Another 5 subclusters demonstrated the most marked transcript enrichment or impoverishment at the decline stage, the last stage of brooding before death (Supplement Fig 4). Because of sample scarcity (only 2 RNA-sequencing samples in decline stage and a limited ability to pursue follow-up studies), we did not explore these subclusters further. Five additional clusters showed monotonic changes and were studied with BLASTP as described below (Supplement Fig 5). No candidate signaling molecules were identified. In total, we excluded 855 genes in 22 subclusters from further analyses.

We focused on the 343 remaining genes in 3 subclusters. Among these subclusters (Fig 4), we saw dramatic transitions in differential gene abundance across behavioral stages. Two subclusters showed increases in expression over time (Fig 4A and B), while the third showed a drop in expression at the non-mated female to feeding mated female transition (Fig 4C).

**Fig 4.**
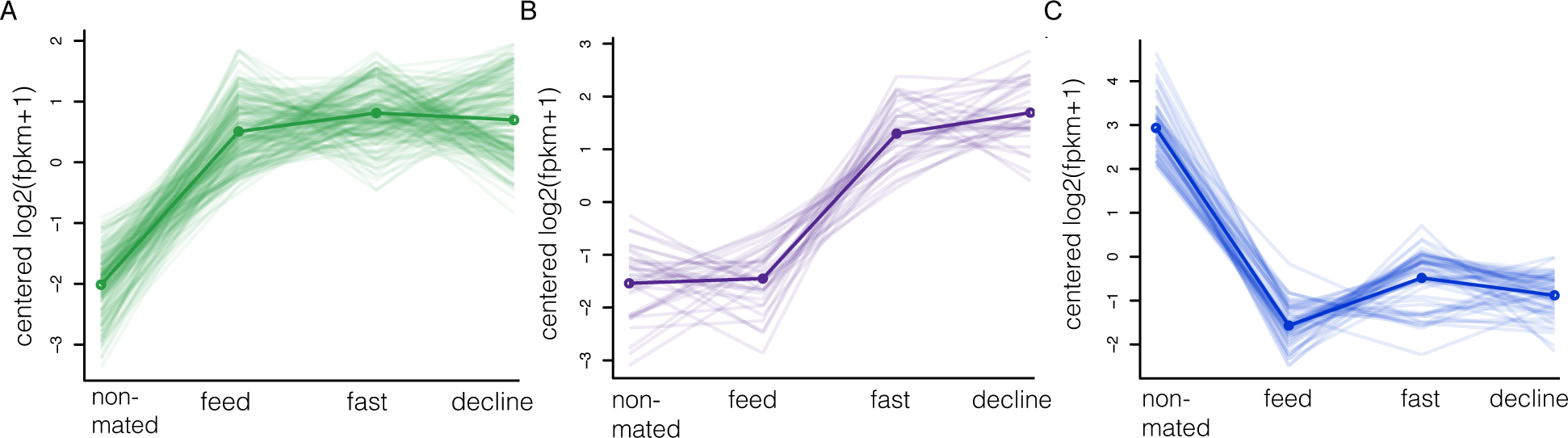
Cohorts of genes are differentially expressed across behavioral stages. Green and purple subclusters identify genes that increased in abundance as the maternal stages progressed (A and B, respectively), while the blue subcluster illustrates genes that decreased over time (C).

**Fig 5.**
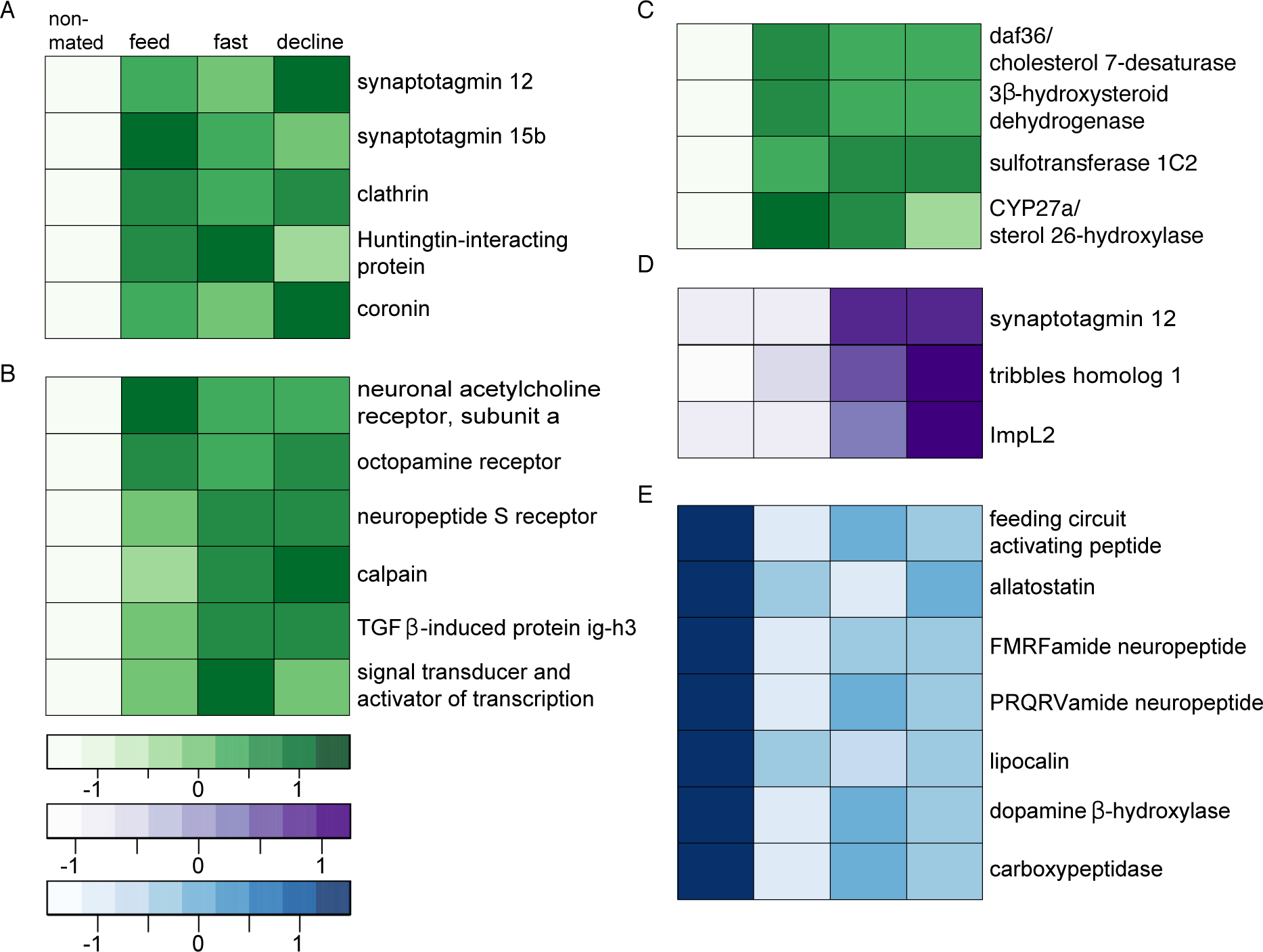
Expression profiles of differentially expressed genes important to optic gland signaling. Heatmap colors correspond to cluster colors depicted in Fig 4. Columns are organized as designated at the top of in panel A. Color key values refer to row z-scores.

We used BLASTN, BLASTP, and PFAM (Altschup et al., 1990; Finn et al., 2016) to identify transcripts (Fig 5). In all clusters examined, there were many housekeeping genes, such as actin and collagen. We also found predicted proteins with no homology to genes of other species in available databases (downloaded October 28, 2015). These uncharacterized genes may represent octopus- or cephalopod-specific proteins. All three clusters did, however, contain a number of transcripts of particular relevance to optic gland biology.

#### 4.2.1 Green cluster Fig 4A: genes in this cluster increased in abundance after mating and remain elevated through the maternal stages

Although the optic gland is regarded as a neuroendocrine organ, its mechanisms of secretory release are unknown. We found transcripts implicated in neural signaling that increased in abundance after mating and remained elevated through all brooding stages (Fig 4A and 5A): synaptotagmin 12, synaptotagmin 15b, and clathrin. Huntingtin-interacting protein (HIP1) and coronin showed a similar expression profile. Although not included in classical models of synaptic transmission, proteins encoded by these two transcripts have in recent years been found to be important for aspects of neuronal signaling (Borlido et al., 2009; Suo et al., 2013). We also saw increases in selected neurotransmitter receptors (Fig 5B), including an alpha subunit of nicotinic acetylcholine receptor, and G-coupled protein receptors similar to octopamine and neuropeptide S receptors in other species. Finally, signaling molecules, including candidate signaling peptides and enzymes implicated in steroid biogenesis, were present in this subcluster (Fig 5C). These results suggest that the optic gland is a site of active signaling during the maternal period, and that the nature of this signaling is different from that of unmated female optic glands.

#### 4.2.2 Purple cluster (Fig 4B): genes in this cluster increased with the transition from feeding to fasting

In this subcluster, we found that the insulin-binding protein Impl2 is enriched in the optic glands of fasting and decline animals (Fig 4B and 5D). This transcript is joined in expression dynamics by two others, tribbles homolog 1 (TRB1) and an additional isoform of synaptotagmin.

#### 4.2.3 Blue cluster (Fig 4C): non-mated optic glands are enriched in neuropeptides that precipitously drop in abundance in brooding females

As in the other subclusters, we found genes encoding proteins involved in cell signaling (Fig 5E), such as lipocalin and dopamine beta-hydroxylase. Particularly notable in this subcluster are the feeding circuit-related neuropeptides and carboxypeptidase (Fig 5E).

#### 4.3 Are molecular changes of the optic glands a reflection of global changes in gene expression?

The abundance of genes encoding neuropeptides in the non-mated optic gland and their subsequent depletion in mated females motivated us to investigate whether molecular changes in the optic gland were mirrored in other tissues of the body. Many of these tissues were sampled in the arm transcriptomes, including axial nerve cords, suckers, muscles, chromatophores, and skin. Using BLASTP and TBLASTN, we baited the arm transcriptomes from all behavioral stages with the genes that were significantly enriched in non-mated optic glands (blue cluster, Fig 4C and 5E). We were unable to recover any of the optic gland neuropeptides in the arm transcriptomes.

## 5. Discussion

With their large central nervous systems and wide range of behaviors, coleoid cephalopods have long fascinated neuroscientists. Pioneering work from Wodinsky and Wells and Wells established that the octopus reproductive axis is similar to that of the vertebrate hypothalamic-pituitary-gonadal axis (Wells and Wells, 1975; Wodinsky, 1977). The optic gland, analogous to the pituitary gland, is the fulcrum of the octopus reproductive axis; it is responsible for innate reproductive behaviors and post-reproductive death. In this study, we confirmed that maternal behaviors can be observed in the lab, including, most importantly, a period of feeding while brooding that precedes the fasting that typically defines the maternal period in octopuses (Movie S1) (Hanlon and Messenger, 1998; Rocha et al., 2001).

Our transcriptomic analyses reveal for the first time a molecular portrait of the optic glands at the distinct behavioral stages of adult female octopus life. In particular, we found that reproduction and the onset of fasting trigger major transcriptional activation in the optic glands. Most importantly, multiple signaling systems of the optic glands, including catecholaminergic, peptidergic, and steroidogenic pathways, are implicated in these behaviors.

### 5.1 Optic Gland Signaling Systems

#### 5.1.1 Feeding peptides

Circulating neuropeptides have been extensively studied in molluscan feeding and reproductive systems (Grimmelikhuijzen and Hauser, 2012; Jékely, 2013). Neuropeptides can influence feeding behaviors through control of motor circuits or modulation of arousal or satiation (Bechtold and Luckman, 2007). Strikingly, we found octopus homologs of these neuropeptides enriched in the optic glands of non-mated female as compared with those of brooding females: feeding circuit activating peptide (FCAP), allatostatin, FMRFamide, and PRQFVamide. The latter 3 peptides have been shown to inhibit feeding behaviors in a diverse range of animals. In Aplysia, FMRFamide and its related neuropeptides inhibits closure of the radula muscle during feeding (Sossin et al., 1987) and in mice, FMRFamide inhibits feeding in food-deprived individuals (Kavaliers et al., 1985). PRQFVamide decreases the excitability of specific neurons in the buccal ganglion feeding central pattern generator and has an overall inhibitory effect on feeding (Furukawa et al., 2003). Allatostatin inhibits feeding in Drosophila by mediating the balance between adipokinetic hormone and insulin-like peptides (Hentze et al., 2015). FCAP, on the other hand, initiates rhythmic feeding motor programs and may be responsible for food-induced arousal (Sweedler et al., 2002).

These data suggest that signaling from an ensemble of neuropeptides tightly controls the feeding behaviors of non-mated animals. In many animals, mechanisms controlling feeding and reproduction are tightly linked and enables metabolic shifts as the individual transitions from growing and maturing to reproduction and tending to young (O’Dor and Wells, 1978). Drosophila females, for example, modify their feeding preferences to consume more protein when they are about to lay eggs (Ribeiro and Dickson, 2010). The reduction in feeding circuit-related neuropeptides at the non-mated to mated transition strongly suggests that, in the octopus, feeding and reproductive state also interact.

Although neuropeptide expression precipitously drops after reproduction, mated animals continue to feed in the initial stage of brooding. However, feeding behavior between mated and non-mated animals manifested in different ways: mated feeding octopuses captured crabs that were within arm’s distance of their dens rather than leave the den to pursue prey, as non-mated females regularly did. This behavioral switch suggests that different feeding strategies are optimal for non-mated and mated females: the feeding-related peptides may regulate energy expenditure or drive to hunt for food. Our data also raise the possibility that redundant peptidergic pathways, or additional non-peptidergic systems, control feeding behaviors. These systems may be independent of the optic gland.

These neuropeptides may also broadly participate in neurotransmission well beyond feeding. If this is the case in the octopus, peripheral tissues would also express these neuropeptides. We checked if these neuropeptides were present in the arm transcriptomes, which included nervous tissue (axial nerve cord), muscle, skin, chromatophores, and suckers. Expression changes of these genes in the arms would suggest that global changes in neuropeptide signaling accompany the aging and senescence process in general. However, all feeding-related neuropeptides in the blue subcluster (Fig 4C and 5E) were missing from the arm transcriptomes, suggesting that their expression is a specific feature of the optic gland. These data are surprising since the optic glands previously have not been implicated in feeding *per se*. Future experiments will reveal if these neuropeptides are functionally conserved in octopuses.

#### 5.1.2 Steroid biosynthesis

Steroids are one of the most evolutionary ancient classes of signaling molecules (Baker, 2011; Nelson and Studer, 2015). Derived from cholesterol, steroids include the sex hormones and corticosteroids. We found that a collection of cholesterol-metabolizing enzymes increased in expression in the first stage of brooding and remained elevated through the maternal period. These included enzymes known to synthesize steroid hormones in diverse animals, from ecdysozoa to deuterostomes (Antebi, 2015). Dafachronic acid in *C. elegans*, for example, regulates longevity and dauer formation, and its biosynthesis relies on the sequestration of cholesterol by daf36/cholesterol 7-desaturase (Yoshiyama-Yanagawa et al., 2011). 3beta-hydroxysteroid dehydrogenase is an oxidoreductase that catalyzes the formation of progesterone in the adrenal gland and gonads of vertebrates (Aakvaag, 1970; Conley and Bird, 1997). Sterol 27-hydroxylase (CYP27a) breaks down cholesterol for both bile acid synthesis and hormone production (Dubrac et al., 2004). Sulfotransferase 1C2 catalyzes the conjugation of a sulfate group to steroid hormones (Foster and Mueller, 2018).

Our study is the first to implicate the optic glands in the production of progesterone and other steroid hormones. Cellular and molecular mechanisms of steroid hormone signaling in octopuses may be different from what has been described previously in the vertebrate neuroendocrine systems. Substrate specificity of these enzymes may differ between species (Antebi, 2015). Functional characterization of these enzymes or identification of steroid metabolites will be particularly useful going forward. Moreover, steroid signaling in molluscs may not be mediated through the receptors orthologous to those identified in vertebrates (Thornton et al., 2003). The *Octopus vulgaris* ortholog of the vertebrate estrogen nuclear hormone receptor, for example, does not bind estrogen (Keay et al., 2006). Our identification of the coordinate enrichment of cholesterol-metabolizing enzymes in mated female octopuses raises the exciting possibility that the optic glands produce a suite of steroid-derived hormones that are specific to the reproductive adult and may have novel cellular targets.

#### 5.1.3 Insulin signaling

Our data reveal that expression of ImpL2 increased at the transition from feeding to fasting (Fig 4B and 5D). In *Drosophila*, ImpL2 is a homolog of insulin-like growth factor binding protein (IGFBP) that reduces activity of the insulin/IGF signaling pathway (Figueroa-Clarevega and Bilder, 2015). The insulin/IGF signaling pathway controls many biological processes, including glycogen metabolism and organismal growth. Insulin-like signaling molecules have been found in many invertebrates (Ahn et al., 2017; Veenstra, 2010), but functional characterization of these molecules in molluscs has been limited to a few species (Gricourt et al., 2003; Hamano et al., 2005). In the Pacific oyster *Crassostrea gigas*, insulin-like molecules stimulate protein synthesis in mantle edge cells, promoting growth in the shell and soft tissues (Gricourt et al., 2003). Seasonal fluctuations in insulin-related peptide expression coincide with body growth and gametogenesis (Hamano et al., 2005).

The actions of the insulin/IGF signaling pathway in octopus are unknown. Females in the decline stage have the highest expression of ImpL2. At this terminal stage of brooding, female octopuses show several indications of wasting, including loss of muscle tone and color. *Drosphila* implanted with malignant tumors secrete high levels of ImpL2 and show marked degradation of tissues, including gonadal, muscle, and adipose tissue (Figueroa-Clarevega and Bilder, 2015; Kwon et al., 2015). These remarkable similarities lead us to predict that the role of Impl2 as an antagonist of the insulin/IGF-signaling pathway is functionally conserved between octopuses and *Drosophila*.

ImpL2-induced tissue wasting is independent of food intake in *Drosophila*: tumor-bearing hosts with high ImpL2 signaling show no differences in food ingestion or feeding behavior from that of control hosts, and high calorie food was insufficient to prevent tissue wasting (Figueroa-Clarevega and Bilder, 2015). In contrast, octopus ImpL2 expression coincides with cessation of feeding. Remarkably, results of high ImpL2 in both *Drosophila* and octopus are consistent with symptoms of cancer cachexia in humans. Patients with cachexia experience an involuntary loss of body mass, undergo substantial degradation of muscle and fat, and report a loss of appetite (Argilés et al., 2014). Medical interventions focusing on increasing caloric intake fail to counteract wasting (Fearon et al., 2012; Op den Kamp et al., 2009)}.

Our results suggest that ImpL2 is intimately tied to feeding behaviors in octopus and implicate the optic glands as a regulator of insulin signaling. These data raise the possibility of a conservation of function in insulin-signaling pathways with animals as distantly related as humans and octopuses. Indeed, clinical work on humans has shown that cachexia and other metabolic conditions are associated with elevated levels of IGFBPs (Clemmons et al., 1991; Huang et al., 2016). Our study, however, is the first direct implication of IGFBP enrichment in a physiological, non-pathological, process.

#### 5.1.4 Neurotransmitter biosynthesis and neurotransmission

The optic gland regulates sexual maturation and behavior by hormone release, but cellular mechanisms of secretion are unknown. We identified a number of genes implicated in neurotransmission and secretion enriched after reproduction (Fig 4A and 5A). Clathrin, for example, facilitates the formation of coated secretory vesicles, and synaptotagmins contribute to the cellular complex that enables vesicle fusion for regulated exocytosis (Brunger et al., 2018; Wu et al., 2014). Specific isoforms of both are enriched in the optic glands at all mated stages. Our finding mirrors existing data on the organization of the vertebrate reproductive axis: synaptotagmins and other proteins involved in classical neurotransmission are differentially expressed in the anterior pituitary gland where they are involved in the release of signaling vesicles and other secretory granules (Jacobsson and Meister, 1996). Moreover, the enrichment of neurotransmitter receptors (Fig 5B), including octopamine receptor, neuropeptide S receptor, and a subunit of nicotinic acetylcholine receptor, suggests that the optic gland undergoes shifts in the signals it receives from the central brain, or in auto- or endocrine signaling, after reproduction.

Optic glands of non-mated animals showed elevated expression of dopamine β-hydroxylase (DBH, Fig 5E), an enzyme involved in the synthesis of catecholamine neurotransmitters. DBH has been shown to catalyze the ß-hydroxylation of dopamine to norepinephrine in vertebrates and tyramine to octopamine in invertebrates (Fernstrom and Fernstrom, 2007; Monastirioti, 1999). These small molecules are used for signaling throughout nervous systems. In the octopus optic gland, candidate aminergic secretory vesicles have been found (Nishioka et al., 1970). Our results suggest that the optic gland engages in endocrine catecholamine signaling and that reproduction leads to the specific shutting down of catecholamine pathways, as opposed to an overall decrease in neurotransmission or cell signaling in the optic glands.

### 5.2 Octopus semelparity

Mechanisms of death differ among animal systems exhibiting semelparity. In several vertebrate species, it is believed that the metabolic cost of gamete production and finding a mate are so high that death is inevitable (Fisher and Blomberg, 2011; Kindsvater et al., 2016). Semelparity in octopuses cannot be explained by the large metabolic output required to produce and deposit eggs: glandectomized animals can reproduce again (Wodinsky, 1977). Instead, it is likely that other evolutionary mechanisms drive post-reproductive death. Coleoid cephalopods are cannibalistic: hatchlings from the same clutch often eat each other, and females sometimes kill males after reproduction (Hanlon and Messenger, 1998). Post-reproductive death is an undeniably effective method in ensuring that the reproductive female does not consume her young. Moreover, longitudinal studies of octopuses, and other molluscs, demonstrate that individuals continue to undergo logarithmic growth as adults (Gricourt et al., 2003; Nixon, 1969; Van Heukelem, 2010; Wells and Wells, 1970). Endless growth without death would so skew octopus populations towards large, old adults that hatchlings would be unlikely to compete. Programmed organismal death in coleoids may exist as a mechanism to safeguard the survival of the next generation of these voracious predators.

In Pacific salmon *Oncorhynchus kisutch*, steroid hormone signaling has been implicated as the cause of death after spawning. Elevated cortisol levels mediate programmed death through tissue degeneration and impaired homeostatic ability (Robertson and Wexler, 1960). Our data show that steroid signaling is one of several pathways implicated in octopus post-reproductive death. Although we do not see molecular evidence of cortisol signaling *per se* in our data, the enrichment of cholesterol-metabolizing enzymes in mated females suggests that steroid metabolites are crucial to semelparity in octopuses. It will be important to identify the steroids in the optic gland and assess the contributions of steroid and ImpL2 signaling, along with the reduction in feeding peptides and catecholamine signaling, in cephalopod post-reproductive death.

### 5.3 Conclusions

Our transcriptome findings suggest that, far from secreting a single hormone, the optic gland likely enlists multiple signaling systems to regulate post-reproductive behaviors and death (Fig 6). Among these are neuropeptidergic systems and cholesterol-derived hormone signaling. Our data raise the possibility that prior to mating, optic gland signaling is dominated by neuropeptides and catecholamines, whereas after mating, it is dominated by steroid hormones.

Wells and Wells originally drew a connection between the direct brain regulation of the octopus optic glands and the vertebrate hypothalamic-pituitary-gonadal axis (Wells and Wells, 1969). Here, we identify a difference between octopus and vertebrate systems: the endocrine signaling of the anterior pituitary gland depend on peptide hormones alone (Davis et al., 2013), whereas our data implicate steroid and catecholamine pathways, in addition to peptide signaling, in the functions of the optic gland. While steroid hormones do affect the actions and feedback of the vertebrate pituitary gland, steroidogenesis occurs in targets of the pituitary gland only, such as the adrenal cortex, and not in the pituitary gland itself (Handa and Weiser, 2014; Werbin and Chaikoff, 1961). Similarly, biosynthesis of catecholamines, such as norepinephrine and dopamine, occurs in the adrenal medulla in response to autonomic stimulation (Fujinaga et al., 1999).

**Fig 6.**
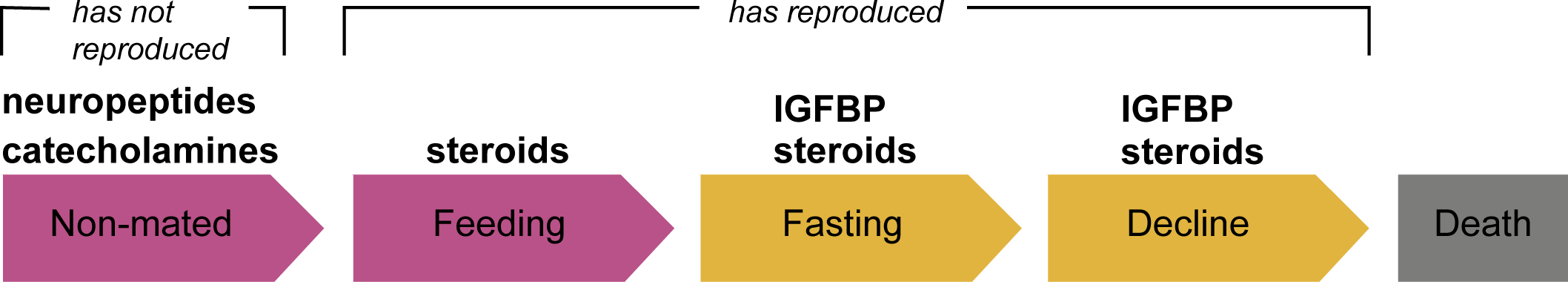
Updated model of optic gland function in maternal behaviors and death. Arrows indicate behavioral groups (pink: feeds; yellow: abstains from feeding). Signaling systems above arrows highlight enriched genes.

If the optic gland encompasses functions of the anterior pituitary gland and the adrenal glands, it prompts the question of what glandular targets of the optic gland, if any, might exist. The oviducal glands, which produce substances that cover the eggs, are candidates. These glands, along with the gonads and oviducts, were found to increase in size as the optic gland enlarges (Wells and Wells, 1959; Wells and Wells, 1975). *In vitro* experiments demonstrated that oviducal secretion is modulated by dopamine and neuropeptides, including FMRFamide (Di Cristo and Di Cosmo, 2007). However, the putative source of direct oviducal control is the fusiform ganglion, not the optic glands. The fusiform ganglion is thought to innervate the oviducal gland, the oviducts, and the systemic heart (Young, 1971). Notably, the fusiform ganglion has been found to contain FMRFamide-like immunoreactive nerve endings (Di Cristo et al., 2003). The optic gland may exert indirect control over glandular targets by signaling to peripheral ganglia, or act in parallel with peripheral ganglia to regulate reproductive tissues.

Our study extends the optic gland-pituitary analogy and uncovers similarities between the optic glands and the vertebrate adrenal glands, demonstrating that the organization and function of the optic glands may be even more pivotal for octopus organismal physiology than previously appreciated.

## 6. Acknowledgements

We are extremely grateful to junior research assistants Emily Garcia, Olivia Harden, Sofia Collins, and Magdalena Glotzer for their contributions throughout the project. We thank Dr. Chuck Winkler and the team at Northeast Brine Shrimp for supplying us with animals. We also acknowledge the UChicago Genomics Facility for library preparation and sequencing, and Mike Jarsulic and the UChicago Center for Research Informatics for indispensable expertise on programming and data management. This research was conducted on the land of the Potawatomi, Miami, Sauk, and Fox peoples.

## 7. Declaration of interests

The authors declare no competing interests.

## 8. Funding

This work was funded by the NSF (IOS-1354898 to CWR, DGE-0903637 to ZYW) and the Erma Smith Endowment. The University of Chicago Genomics Facility is supported by the NIH (UL1 TR000430).

